# Mistakes in the re-analysis of lipidomic data obtained from a human model of resolving inflammation lead to erroneous conclusions

**DOI:** 10.1101/2023.03.14.532368

**Authors:** Jesmond Dalli

## Abstract

Recent years have seen an increased interest in the biology of specialized pro-resolving lipid mediators (SPM) with many investigators evaluating both their endogenous production as well as their biological and pharmacological properties. This increased interest has led to a rapid evolution in our understanding of both the biological and pharmacological activities of these mediators with their endogenous formation and biological activities being documented in a wide range of species that spans the evolutionary tree including fish, planaria and humans. Despite this plethora of evidence in a recent article Homer and colleagues claim that the reanalysis of a published dataset - partly originating from our laboratory - demonstrates that these molecules are absent in human blister exudates, contrary to the evidence published earlier. Based on their observations these Authors conclude that SPM are not relevant to human resolution biology. In this manuscript a reanalysed the dataset using unbiased methodologies and criteria that align with those recently proposed by the UK Consortium on Metabolic Phenotyping was performed together with the re-analysis of the dataset using criteria described by Homer and colleagues. Results from this re-analysis confirmed presence of SPM in human blister exudates and that the methodologies employed for quantitation of these molecules are robust. We also discuss how the results obtained in the article published by Homer and colleagues present several points of concern including the use of employ an arbitrary cut-off value to assign the noise for all the transitions used that does not take into consideration the fluctuation of the signal in each transition and therefore is not truly representative of the background signal. The use of different transitions to those employed in the original analyses and misreporting of findings based on the criteria employed. In conclusion the evidence presented herein demonstrates that correct application of rigorous criteria accepted by the community is essential in ensuring accurate identification of mediators and avoid blatant mistakes which can impact on the scientific development of the field.

## Introduction

Understanding mechanisms that regulate host immunity to terminate inflammation and facilitate tissue repair is essential to gain better insights into both disease mechanisms and the development of novel therapeutics for the treatment of these conditions. There is now extensive evidence supporting a central role for specialized pro-resolving mediators (SPM) in regulating these two fundamental mechanisms (i.e. inflammation resolution and tissue repair). Recent studies from many independent investigators have linked alterations in the endogenous production of these molecules with disease propagation and severity ^1–4^ as well as with the protective activities of omega-3 supplements ^5–7^. Studies by Gilroy and colleagues have indicated how efficient engagement of SPM, namely 15-epi-lipoxin A4, links with the ability of humans to resolve inflammation as well as to respond to the anti-inflammatory activities of aspirin ^8,9^. As part of the efforts to understand the endogenous role that these molecules plan in the regulation of inflammatory responses in vivo, in collaboration with Gilroy and other colleagues in 2016 we evaluated production of SPM and other lipid mediators in blister exudates ^10^. In the original study, we reported the temporal regulation of three SPM, Resolvin (Rv) D5, Lipoxin (LX) B4 and RvE3 in blister exudates. We also identified several SPM that were present in a subset of the samples analysed. More recently, Homer and colleagues have sought to re-analyse this original dataset employing distinct identification criteria to determine the robustness of the original observations ^11^. This additional scrutiny is welcome since it allows for an open discussion around the endogenous role of SPM in humans. However, in reviewing the pre-print of the manuscript it became apparent that several mistakes were made in the analytical methodology that might have led to erroneous results and conclusions. Supported by more broadly accepted analytical approaches, herein a detailed description of these errors is discussed along with a re-analysis of the data employing rigorous and widely used standards that support the original findings.

## Methods

Blister samples reanalysed in the present manuscript were collected from the timepoints:

- A1, A2, A3, A4 (These samples were collected at the 48h interval)
- B1 (This sample was collected on Day14)
- C1, C2, C3 (These samples were collected at the Day 17 time interval)
- D1, D2 (Sample D1 was collected at the 8 h interval and D2 was collected at the 14 h interval)

In addition to these samples we the dataset included mixes that contained the relevant synthetic or authentic standards for the following mediators:

RvD1; RvD2; RvD3; RvD4; RvD5; RvD6; 17-RvD1; 17-RvD3; PD1; 10S, 17SdiHDHA; MaR1; 7S, 14S-diHDHA; LXB4; RvE3, 5S,15S-diHETE; RvE1; RvE2; LXA4; 15-epi-LXA4

### Reanalysis criteria

#### Replication of analysis performed by Homer and colleagues

In the first instance we sought to replicate the analysis performed by Homer and colleagues using the criteria described in their manuscript. Here we employed a signal of 3000 cps as the cut-off for the lower limits of quantitation and a signal of 10000 as a cut-off for the lower limits of detection.

#### Reanalysis of dataset using unbiased criteria

In this reanalysis we defined the lower limits of detection as a peak with a signal to noise ratio of 3:1 whereas the lower limits was of quantitation were defined as a signal to noise ratio of 5:1 in accord with criteria proposed by UK Consortium on Metabolic Phenotyping and in line with guidelines from international bodies (see ^12^). Signal to noise ratios were calculated using the Relative Noise algorithm using Sciex OS 3.0. Further evidence for the identity of the mediators of interest was obtained by matching the MS/MS spectra obtained with those from authentic standards for each of the mediators using the library match function in Sciex OS 3.0. A match score ≥ 70% was used as a cut-off for positive match. Standard curves were prepared using Sciex OS 3.0 and authentic standards for the mediators of interest using a linear regression analysis and weighting factor of 1/x^2^. The relative mean error for points on this curve was then calculated to determine the accuracy of the curve with a cutoff value of <20%.

## Results

### Omission of data that fulfils stipulated criteria

In their methodology Homer and colleagues state that they used a 3000 cps cut-off as their limit of detection i.e. the limit that determines whether a specific molecule is present or absent in a given sample. Re-analysis of the dataset demonstrates that the following mediators: RvD4, LXA4, LXB4 and RvD6 are clearly above this cut-off (Figure 1), nonetheless the Authors state in the manuscript that these molecules were not identified in the samples.

**Figure 1:**
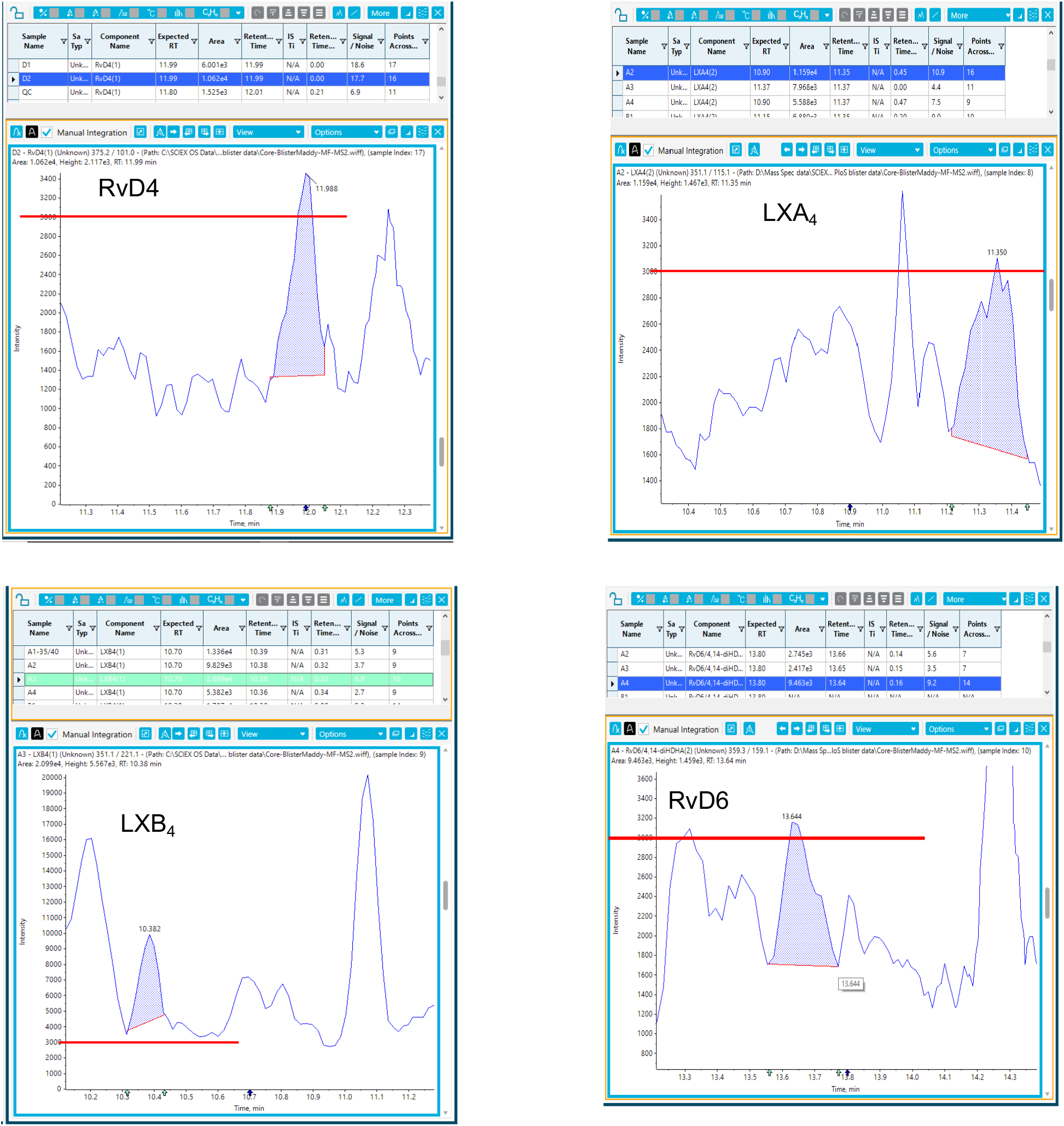
Figure highlighting exemplar peaks where the signal is greater than 3000 cps employed as the lower limits of detection. Red line denotes the 3000 cps cut-off.

The Authors also state that they use a cut-off of 10000 cps as their limit of quantitation and claim that both RvD5 and LXB4 did not meet this criterion. However, a re-analysis of the data demonstrates that signals obtained for these two mediators are both above this arbitrary cut-off value (Figure 2).

**Figure 2:**
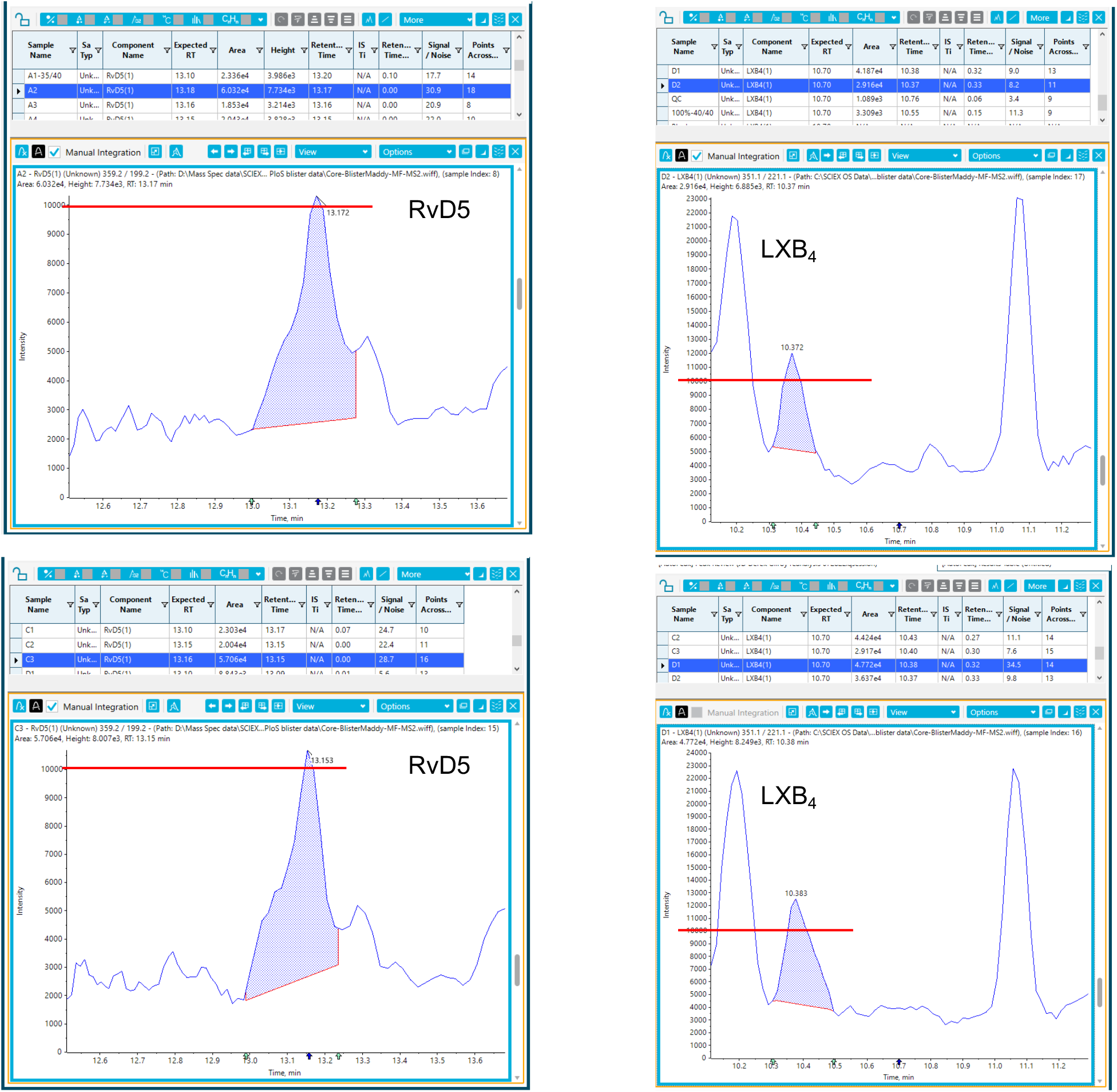
Figure highlighting exemplar peaks where the signal is greater than 10000 cps employed as the lower limits of quantitation. Red line denotes the 10000 cps cut-off.

Together these findings indicate that the conclusion drawn by the Homer and colleagues are not supported by the results they presented.

### Use of rigorous criteria corroborates the presence of SPM in blister exudates

To extend the discussion on the robustness of the data presented in the original article a reanalysis was performed using rigorous criteria used in the field and, more recently, recommended by the UK Consortium on Metabolic Phenotyping, which is in line with guidelines from international bodies (see ^12^). Here using unbiased methodologies, a signal to noise ratio of 3:1 was employed to define the lower limits of detection and a signal to noise ration of 5:1 was employed to define the lower limits of quantitation. This approach demonstrates that all the SPM reported in Motwani et al were identified (i.e. a signal to noise ratio ≥ 3:1) in blister samples (Figure 3). The identity of the SPM was further corroborated by evaluating MS/MS spectra and comparing these to those obtained using reference standards (Figure 4). Furthermore, the SPM quantified in the published article (see Figure 3 in^10^), all gave a signal that was greater to the lower limits of quantitation (Figure 3).

**Figure 3:**
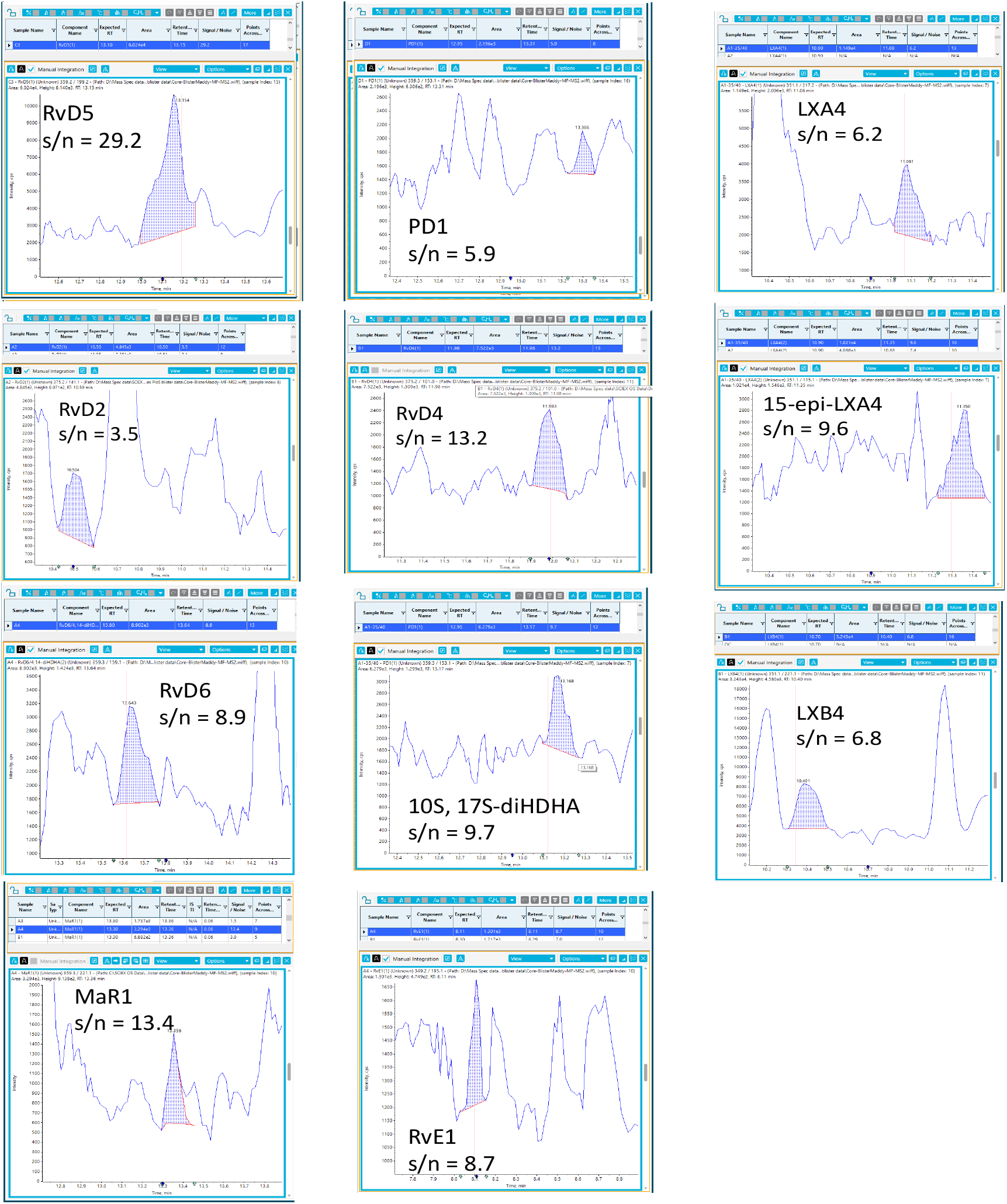
Evidence supporting the presence of SPM in blister exudates.

**Figure 4:**
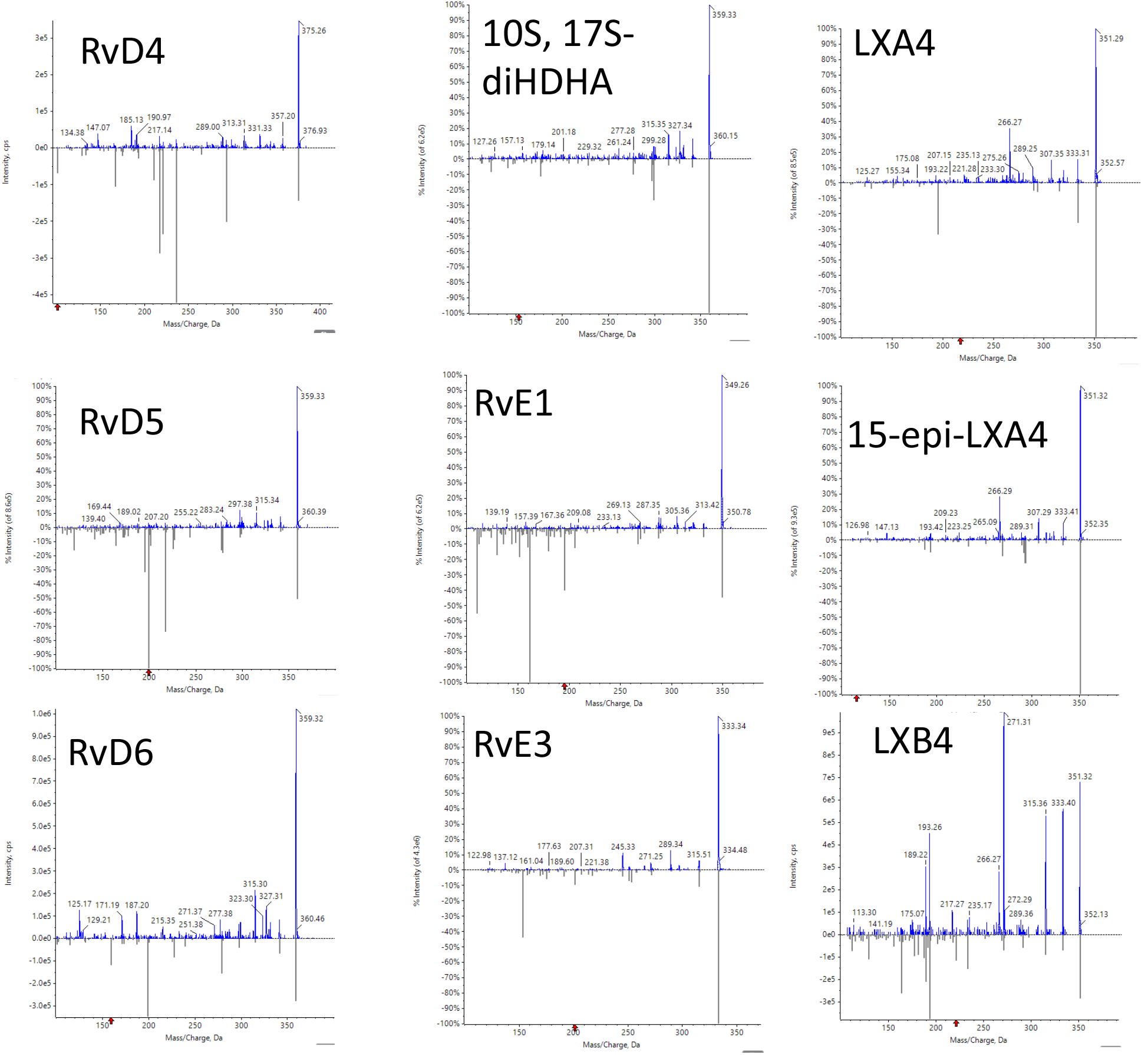
MS/MS spectra employed in the identification of SPM in blister exudates. MS/MS spectrum in blue is obtained from blister exudates, whereas MS/MS spectrum in grey is the reference spectrum.

### Use of weighted standard curves demonstrates accuracy of std curves

It is accepted that LC-MS/MS datasets are heteroskedastic and, as such, the use of weighting is widely accepted to ensure that the results obtained from such standard curves are accurate and precise. Indeed, when we applied weighted standard curves for mediators quantified in Motwani et al.^10^, we observed that the relative error of the mean was < then the 20% cut-off value recommended, ranging between 0.3 and 13.1%. Therefore, we are confident that quantifications presented in the original publication are robust.

## Discussion

The present manuscript demonstrates the importance of utilising unbiased methodologies in the analysis of lipid mediators. Key aspects that need to be taken into consideration in such analysis include the identification of the noise threshold to establish whether the signal obtained at the retention time that corresponds with that of the mediator of interest is a critical step. Distinct approaches are employed in the literature for this task with the main criterion of the various methods employed being to determine the fluctuations in the background signal to determine whether the signal (peak) of interest is distinct from the background noise at a level that can be employed to reliable identify and quantify the mediator (https://www.agilent.com/cs/library/technicaloverviews/public/5990-7651EN.pdf).

In their re-analysis, the Homer *et al* state that’ *Baseline was assigned adjacent to, and ahead of the peak of interest in the biological samples. The baseline was generously assigned as 1,000 peak height broadly across the dataset’*. However, scrutiny of the dataset demonstrates that the signal in the regions noted by the Authors widely varies across the dataset as noted in Figure 5. Furthermore, such an approach does not take into consideration the variations across the baseline which constitutes the noise to be accounted for. Thus, using such an approach will lead to both over and under-estimation of the noise component within the chromatogram, leading to erroneous results and conclusions.

**Figure 5:**
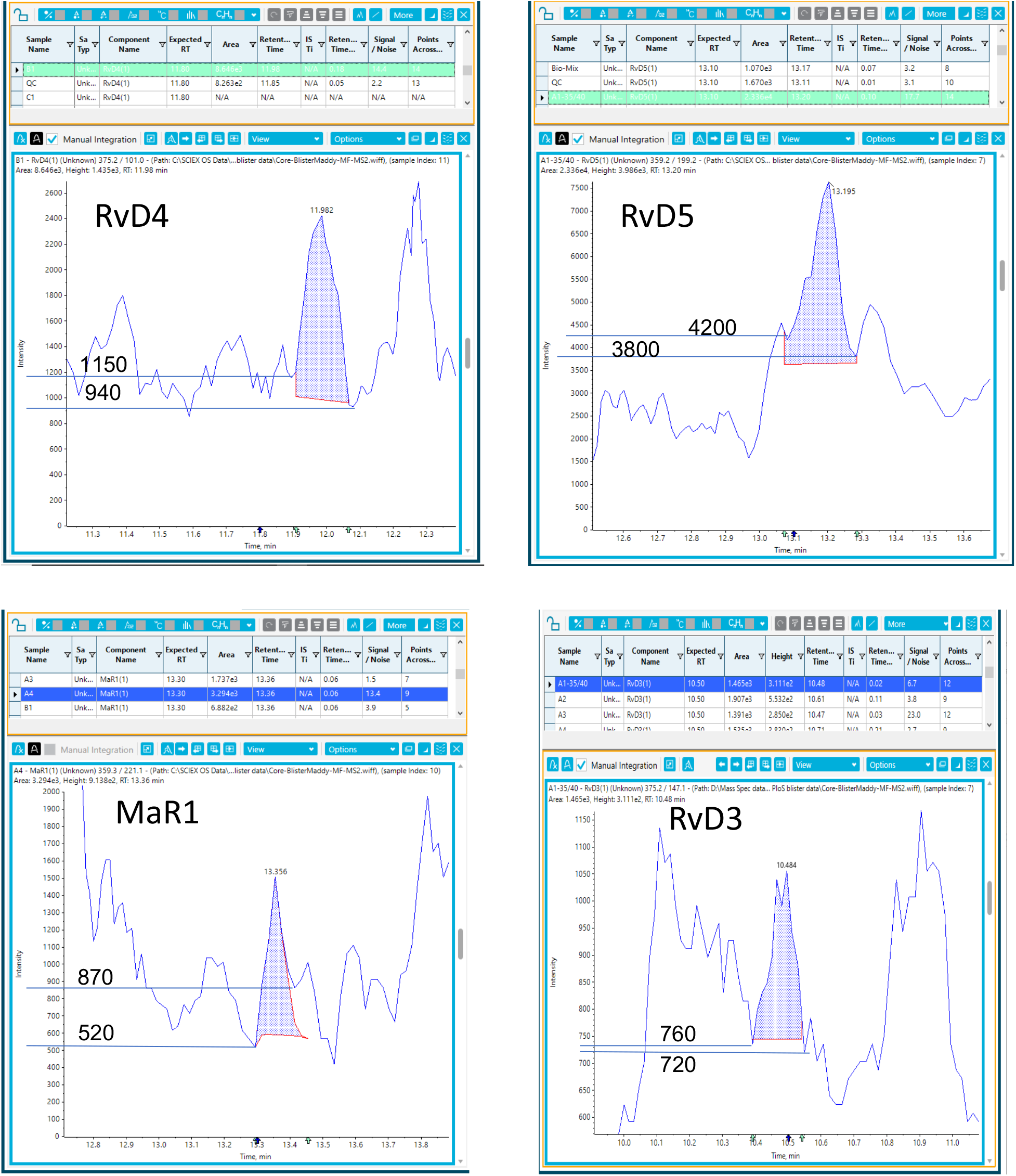
Figure highlighting variations in the baseline signal immediately before and after the peak of interest.

It is also essential that when datasets are re-analysed with the intent data replication, the same Multiple Reaction Monitoring (MRM) transitions are employed to those employed in the initial analysis^10^. This is because each of these transitions is likely to yield distinct matrix response signals and the abundance of the signal obtained for each ion pair is likely to be different. In their reanalysis Homer and colleagues employed different transitions to those employed in the original analysis for some of the mediators, namely for: MaR1, 7S, 14S-diHDHA, 17R-RvD3, 17R, RvD1, 10S, 17S-diHDHA, RvE1 and RvE2. The information related to the transitions employed in the analysis of this subset of samples was made available to the co-authors both in 2016 and, again, in August 2022. For independent scrutiny, we provide the tables provided to the co-authors on these two occasions (Table 1 and Table 2).

**Table 1:** Table provided in August 2016 to Prof Gilroy for data presented in^10^.

|  | MRM transitions |  |
| --- | --- | --- |
|  | Q1 | Q3 |
| RvD1 | 375 | 141 |
| RvD2 | 375 | 141 |
| RvD3 | 375 | 137 |
| RvD4 | 375 | 101 |
| RvD5 | 359 | 199 |
| RvD6 | 359 | 159 |
| 17R-RvD1 | 375 | 233 |
| 17R-RvD3 | 375 | 147 |
| PD1 | 359 | 153 |
| 10S,17S-diHDHA | 359 | 153 |
| MaR1 | 359 | 221 |
| 7S,14S-diHDHA | 359 | 221 |
|  | Q1 | Q3 |
| RvE1 | 349 | 195 |
| RvE2 | 333 | 199 |
| RvE3 | 333 | 201 |
|  | Q1 | Q3 |
| LXA<sub>4</sub> | 351 | 115 |
| LXB<sub>4</sub> | 351 | 221 |
| 5S,15S-diHETE | 335 | 235 |
| 15R-LXA<sub>4</sub> | 351 | 115 |
| 15R-LXB<sub>4</sub> | 351 | 221 |
| LTB<sub>4</sub> | 335 | 195 |
| 20-OH-LTB<sub>4</sub> | 351 | 195 |
| PGD<sub>2</sub> | 351 | 189 |
| PGE<sub>2</sub> | 351 | 189 |
| PGF<sub>2α</sub> | 353 | 193 |
| TxB<sub>2</sub> | 369 | 169 |

**Table 2:**
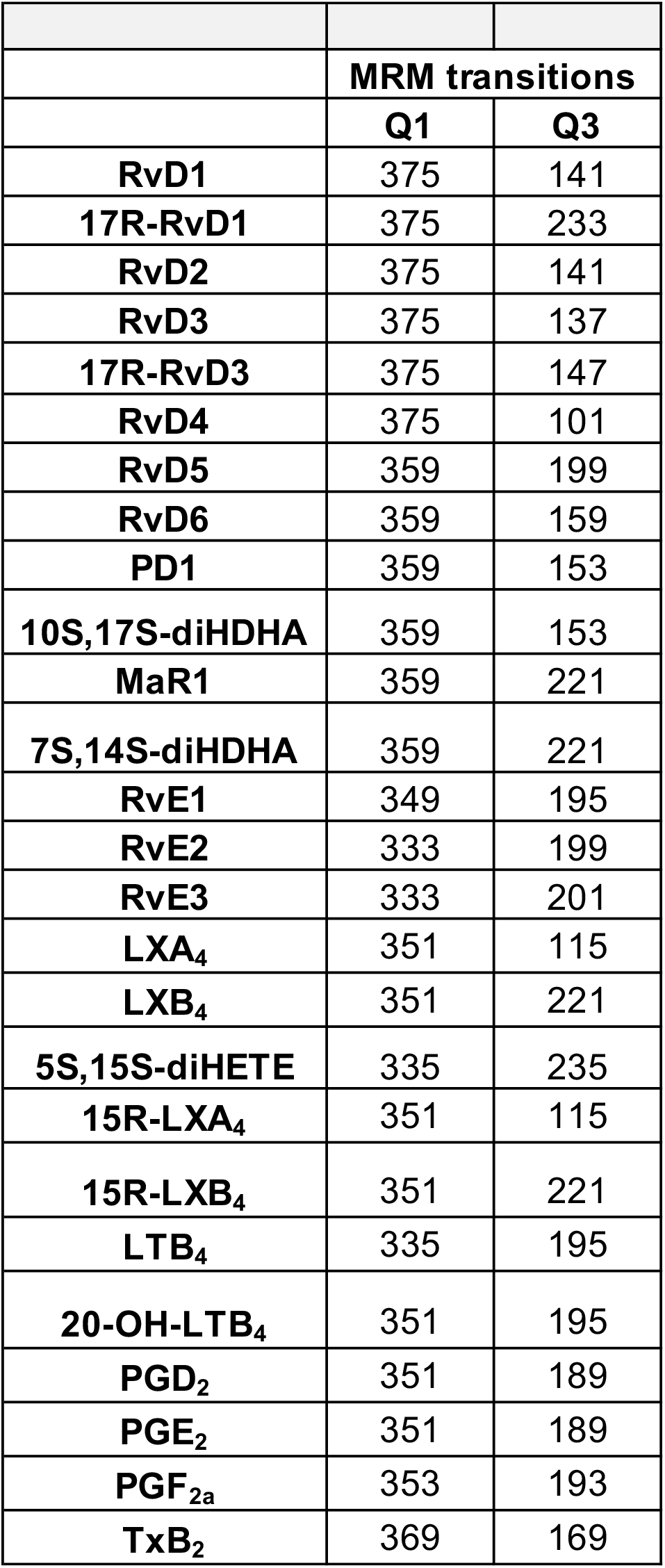
Table provided to Homer and colleagues in August 2022 for data presented in ^10^.

Finally, the application of cutoff values is a critical issue that has been widely discussed in the field of metabolomics with various efforts to obtain consensus criteria cross the field to facilitate the development of both basic research and biomarker discovery^12^. In their re-analysis of Homer and colleagues state that community accepted criteria for limits of detection and limits of quantitation are of a signal to noise ratio of 3:1 and 10:1, respectively. Whilst they are correct that many investigators use a signal to noise ratio of 3:1 as the limits of detection, a signal to noise ratio of 10:1 is not a widely accepted criterion in the field. The latter aspect is reflected in several recently published guidelines including, most recently, the one provided by the UK Consortium on Metabolic Phenotyping: herein, it is proposed that a signal to noise ration of 5:1 as the cut-off for determining limits of quantitation ^12^.

Therefore, in conclusion evidence presented herein lends further support to the identification of SPM in human blister fluids, in line with published results. Furthermore, they underscore the importance of establishing consensus criteria for the identification of bioactive molecules. They also highlight the importance of utilising unbiased approaches for the analysis of lipid mediator datasets since application of arbitrary criteria leads to erroneous results and flawed conclusions, in this manner be of detriment to the field.

